# Drug-target interaction prediction using a multi-modal transformer network demonstrates high generalizability to unseen proteins

**DOI:** 10.1101/2023.08.21.554147

**Authors:** Alexander Kroll, Sahasra Ranjan, Martin J. Lercher

## Abstract

Most drugs are small molecules, with their activities typically arising from interactions with protein targets. Accurate predictions of these interactions could greatly accelerate pharmaceutical research. Current machine learning models designed for this task have a limited ability to generalize beyond the proteins used for training. This limitation is likely due to a lack of information exchange between the protein and the small molecule during the generation of the required numerical representations. Here, we introduce ProSmith, a machine learning framework that employs a multimodal Transformer Network to simultaneously process protein amino acid sequences and small molecule strings in the same input. This approach facilitates the exchange of all relevant information between the two types of molecules during the computation of their numerical representations, allowing the model to account for their structural and functional interactions. Our final model combines gradient boosting predictions based on the resulting multimodal Transformer Network with independent predictions based on separate deep learning representations of the proteins and small molecules. The resulting predictions outperform all previous models for predicting drug-target interactions, and the model demonstrates unprecedented generalization capabilities to unseen proteins. We further show that the superior performance of ProSmith is not limited to drug-target interaction predictions, but also leads to improvements in other protein-small molecule interaction prediction tasks, the prediction of Michaelis constants *K*_M_ of enzyme-substrate pairs and the identification of potential substrates for enzymes. The Python code provided can be used to easily implement and improve machine learning predictions of interactions between proteins and arbitrary drug candidates or other small molecules.

## Introduction

The prediction of protein-small molecule interactions is a long-standing challenge in the discovery of new drugs and the understanding of their action^1–11^. The main obstacle to achieving such predictions lies in generating effective numerical representations of the two molecule types that encode all the information relevant to the underlying task. Ideally, these numerical representations should already incorporate information about the molecular and functional interactions between proteins and small molecules. Some recent models have attempted to address this challenge by facilitating the exchange of information between the small molecule representation and the protein representation during their generation^2,3,7^. While these approaches offer the potential to better capture the interplay between the two modalities, they still cannot capture the full complexity of protein-small molecule interactions.

Typically, protein information is incorporated into the small molecule representation only after transforming the protein sequence into a single numerical vector, such that no detailed information on individual amino acids is provided. Similarly, small molecule information is presented to the protein representation only after transforming the small molecule information into a single numerical vector that summarizes the properties of individual atoms and their relationships across the whole molecule. This approach results in the loss of information relevant to the downstream prediction task. Instead, one should consider the entirety of both protein and small molecule simultaneously during the creation of their numerical representations. The resulting representation of a protein could be trained to incorporate information specific to small molecules it interacts with according to the training data, while the small molecule representation could integrate information on its protein partners. However, achieving this detailed information exchange poses a challenge, as proteins and small molecules are represented using different modalities. While proteins are commonly represented by their amino acid sequences, small molecules are represented in much greater detail, often as strings containing information about every atom and bond in the molecule^12^. Consequently, separate deep learning models are typically employed for each molecule type, obstructing effective information exchange.

Popular models for the representation of proteins^13^ and small molecules^14^ are Transformer Networks, which were originally developed for Natural Language Processing (NLP) tasks. Until recently, Transformer Networks focused primarily on a single input modality, such as text or images. However, the past three years saw the development of multimodal Transformer Networks that project two different modalities onto the same embedding space, facilitating the processing of data that integrates both types of information^15–18^. Notably, such multimodal approaches have achieved significant performance improvements over state-of-the-art methods in tasks involving combinations of images and text.

Inspired by this progress, we here present a novel approach to the prediction of protein-small molecule interactions. Our ProSmith (PROtein-Small Molecule InTeraction Holistic) model leverages the power of a multimodal Transformer Network architecture to process protein amino acid sequences and small molecule strings within the same input sequence. For final model predictions, ProSmith combines predictions from three gradient boosting models that utilize (i) a learned representation of the multimodal Transformer Network, (ii) separate general representations of the proteins and the small molecules, and (iii) a combination of all three representations. ProSmith outperforms previous models in predicting the affinity between a protein and a drug, with the greatest improvements seen for the generalization to proteins that were not part of the training set. In addition, we show that ProSmith can also be applied successfully to tasks other than drug-target interaction predictions: ProSmith outperforms previous state-of-the-art models for evaluating whether a small molecule is a natural substrate for a given enzyme^19–27^ and for predicting the Michaelis constant *K*_M_ of an enzyme for its substrate^28–32^.

## Results

### Architecture of the multimodal Transformer Network for the prediction of protein-small molecule interactions

As its input, the ProSmith model takes two modalities, a protein amino acid sequence together with a SMILES string that represents the molecular structure of the small molecule. When constructing multimodal Transformer Networks, different architectural designs can be considered, each differing in how information between the modalities is incorporated^33^. This subsection provides a detailed description of the architectural choices made in the development of ProSmith.

We adopt a concatenation approach that combines the protein amino acid sequence and the SMILES string into a single input sequence. This choice allows the exchange of all protein and small molecule information at any update step. To process an input, Transformer Networks divide the input sequence into small chunks, referred to as tokens. For protein sequences, each amino acid is treated as a separate token, while the SMILES strings for small molecules are divided into tokens as described in Ref.^14^. To facilitate efficient training, ProSmith leverages pre-learned token representations from Transformer Networks that were trained independently on each modality. For amino acid representations, we utilize embeddings from the ESM-1b model, a 33-layered Transformer Network that was trained in a self-supervised manner on 27 million protein amino acid sequences^13^. For the SMILES string tokens, we use embeddings from the ChemBERTa2 model, a Transformer Network trained on a dataset of 77 million SMILES strings^14^ (see level A in Figure 1). The token embeddings derived from the ESM-1b model and the ChemBERTa2 model have different dimensions, 1280 and 600, respectively. To process these embeddings using the same Transformer Network, we employ linear layers to map both sets of embeddings to a joint embedding space with a shared dimension of 768, which is the hidden dimension of all tokens in the ProSmith Transformer Network (level B in Figure 1).

**Figure 1.**
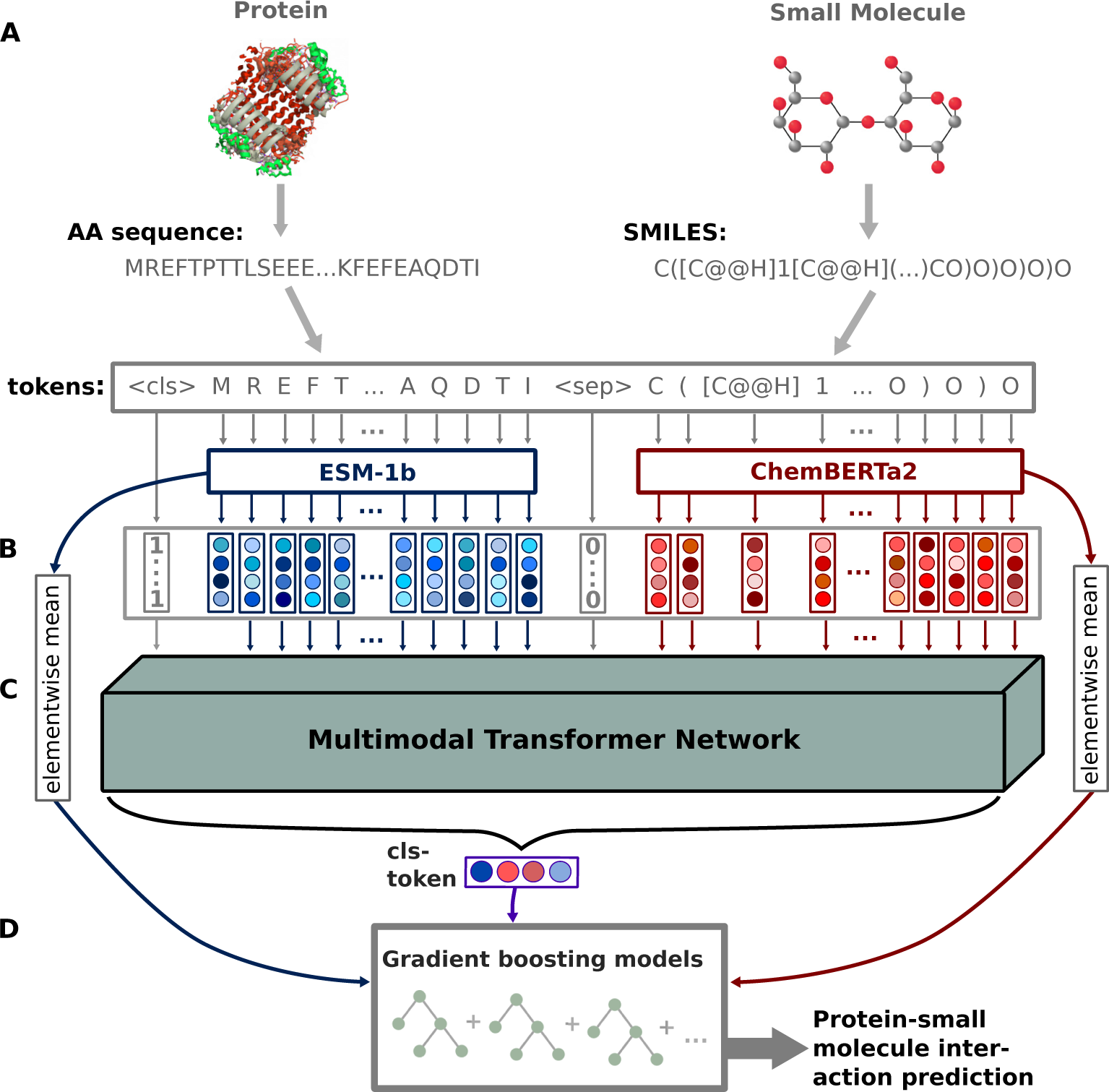
ProSmith model overview. In level A, a protein amino acid sequence and a small molecule SMILES string are transformed into input tokens. The protein tokens are converted to embedding vectors using the trained ESM-1b model, while the SMILES tokens are mapped to embedding vectors using the trained ChemBERTa2 model. In level B, all tokens are mapped to the same embedding space, and are utilized as input sequence for a Transformer Network. In level C, the Transformer Network processes the input tokens and puts out an updated embedding of a classification token (cls), which incorporates information from both the protein and small molecule. In level D, this cls vector, in combination with the ESM-1b and ChemBERTa2 vectors, serves as the input for gradient boosting models trained to predict protein-small molecule interactions.

In addition to the protein sequence and the SMILES string tokens, we add two special tokens: the classification token ‘cls’, its representation is trained as the combined enzyme-small molecule input for downstream tasks, and the separation token ‘sep’, which is identical across input sequences and indicates to the Transformer Network the end of the protein sequence and the start of the SMILES string within the input sequence (level B in Figure 1). During the processing steps, the Transformer Network (level C in Figure 1) updates each input token using the attention mechanism^34^, which enables the model to look at the whole input sequence and to selectively focus only on relevant tokens for making updates. After updating all input tokens for a pre-defined number of steps, the classification token cls is extracted. This token is then used as the input for a fully connected neural network, which is trained to predict an interaction between the small molecule and the protein. By training the entire model end-to-end, ProSmith learns to store all relevant information for the interaction prediction within the cls token.

We set the number of attention layers in the Transformer Network to six, each with six attention heads. In Transformer Networks, the numerical representation of the input sequence is processed in every layer by each attention head separately, updating the input tokens. The updated tokens from all six attention heads are then concatenated and passed as input to subsequent attention layers. The resulting token embeddings from each attention head have a dimension of 128, resulting in the model’s hidden dimension of 768.

### ProSmith feeds the learned representations to gradient boosting models

Following the training of the Transformer Network for predicting interactions between proteins and small molecules, we extract the cls token as a task-specific joint representation for a given protein-small molecule pair. However, due to the limited size of the cls token and the number of update steps, we hypothesized that some relevant general information of the protein and the small molecule might be lost during the generation of the representation. To address this concern, we also more directly use the information contained in the ESM-1b representation of the raw protein sequence and the ChemBERTa2 representation of the SMILES string. We create a single representation for a given protein by calculating the element-wise mean^35^ across its ESM-1b token embeddings and a single representation for a given small molecule by calculating the element-wise mean across its ChemBERTa2 token embeddings. In the following, we refer to these compressed vectors as ESM-1b vector and ChemBERTa2 vector, respectively.

Previous studies have demonstrated benefits of utilizing learned representations from Transformer Networks as inputs for gradient boosting models, leading to improved outcomes compared to directly using the predictions of a Transformer Network^19,36^. We thus follow a similar approach here. Gradient boosting models consist of multiple decision trees that are constructed iteratively during training. In the initial iteration, a single decision tree is built to predict a protein-small molecule interaction of interest for all training data points. By constructing new decision trees, subsequent iterations aim to minimize the errors made by the existing trees. Ultimately, an ensemble of diverse decision trees is formed, each focusing on different aspects of the input features and collectively striving to predict the correct outcome^37,38^. In this study, we leverage the learned cls tokens, ESM-1b vectors, and ChemBERTa2 vectors as inputs for the gradient boosting models.

When aiming to increase model performance, it is a common strategy to train multiple different machine learning models using the same input representations. Using an ensemble of these models, i.e., calculating a (weighted) mean of the various model predictions, can lead to more robust and accurate predictions^39^. We hypothesized that similarly, more robust and improved predictions can be achieved by an ensemble of the same machine learning model trained with different input representations. In a previous study, we indeed observed that training multiple gradient boosting models with different input vectors and combining their predictions through weighted averaging yielded enhanced performance compared to a single model using all input information simultaneously^30^. Thus, we train three distinct gradient boosting models (level D in Figure 1): one using only the cls token, another using the concatenated ESM-1b vector and the ChemBERTa2 vector, and a third model concatenating all three input vectors. To obtain the final prediction, we compute a weighted mean of the predictions from these models, with the weights determined through hyperparameter optimization.

### Model training and hyperparameter optimization

Each dataset used in this study was divided into three subsets used for training, validation, and testing, respectively. The training and validation sets were utilized for hyperparameter optimizations, where different hyperparameter combinations were used to train the model on the training data, and the set of hyperparameters that yielded the best results on the validation set was selected for the final model. The hyperparameters include learning rate, number of hidden layers, hidden dimension, and batch size (a full list is given in Supplementary Table 1).

Due to the substantial time and resource requirements associated with training large Transformer Networks, conducting a systematic hyperparameter search for the multimodal Transformer Network was not feasible on the available hardware. Instead, we employed a trial-and-error approach to identify a suitable set of hyperparameters. We iteratively adjusted the hyperparameters with the aim of improving the results for the drug-target affinity prediction task (see below) on the validation set. The resulting combination of hyperparameters (Supplementary Table 1) was used for all tasks in this study. We trained each Transformer Network for 100 epochs. To guard against overfitting, we performed early stopping, i.e., we saved model parameters after each epoch and finally selected the model that achieved the best performance on the validation set.

For the gradient boosting models, we were able to perform a systematic hyperparameter search to identify the optimal configuration for each task. We conducted random searches^40^ that iterated through 2 000 combinations of hyperparameters, including learning rate, depth of trees, number of iterations, and regularization coefficients (a full list is provided in Supplementary Table 2). After identifying the gradient boosting models that demonstrated the most promising performance on the validation sets, we proceeded to train new models using both the training and validation sets. This final model was then evaluated using the previously untouched test set, ensuring an unbiased assessment of the model’s predictive capabilities and ability to generalize.

### ProSmith outperforms previous models for drug-target affinity predictions

The process of drug discovery is inherently time-consuming and expensive. A crucial aspect of drug dis-covery is the determination of interactions between potential drug compounds, typically small molecules^41^, and their target proteins. Machine learning models that facilitate large-scale predictions of drug-target affinities (DTAs) have the potential to accelerate the overall drug discovery process by identifying appro-priate drug molecules for desired target proteins^42^.

Here, we assess the performance of ProSmith on one of the most widely used datasets for validating drug-target affinity prediction models, the Davis dataset^43^. The Davis dataset comprises 30 056 data points, consisting of binding affinities for pairs of 72 drugs (small molecules) and 442 target proteins, measured as dissociation constants *K_d_* (in units of nM). To create target values for ProSmith, we follow previous prediction methods by using log-transformed values, defined as 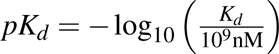. The resulting values range from 5.0 to 10.8. To split the Davis dataset, we adopted the identical strategy employed by the previous state-of-the-art method, NHGNN-DTA by He et al.^2^. He et al. split the Davis dataset into 80% training data, 10% validation data, and 10% test data. Four different scenarios were investigated^2^: (i) a completely random split that includes drugs and targets in the test and validation sets that also occurred in other combinations in the training set (random split); (ii) a split that excludes target proteins used for training from the test and validation sets (cold target); (iii) a split that excludes drugs used for training from the validation and test sets (cold drug); and (iv) a split that excludes any drug and any target that were used for training from the validation and test sets (cold drug & target). To obtain accurate estimates of the true model performance, He et al. created five random splits for each of the four aforementioned scenarios. To ensure a fair comparison between ProSmith and NHGNN-DTA, we followed the same procedure, generating the random splits with the code provided in Ref. [2].

The Davis dataset contains only approximately 30 000 data points, which can be considered relatively small. When the available training data for a specific prediction task is limited, a common strategy in deep learning is to pre-train the model on a related task for which more abundant training data is available^44,45^. To construct such a larger pre-training dataset, we extracted drug-target pairs from BindingDB^46^ with experimentally measured IC50 values; these values indicate the concentration of a drug required to inhibit a target by 50%. We excluded all targets and drugs present in the Davis dataset, thereby ensuring that for the cold splitting scenarios, ProSmith has indeed never seen any of the relevant test targets or test drugs before. The resulting dataset comprised approximately one million drug-target pairs with known IC50 values. We pre-trained the ProSmith Transformer Network for six epochs on this dataset (see Methods for additional details). Subsequently, the learned parameters were used as initial parameters to train the ProSmith Transformer Network on the Davis dataset.

We trained ProSmith on the training and validation data of all five random training-validation-test splits for all four splitting scenarios introduced in Ref. [2]. In the following, we state model performance metrics for each scenario as the mean scores resulting from model validation on the five different test sets. To evaluate model performance, we employ performance metrics that have been used widely in previous DTA prediction studies: the mean squared error (MSE); the concordance index (CI); and the 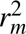 metric. The CI assesses the ability of a predictive model to correctly rank pairs based on their predicted values. It is defined as the fraction of correctly ordered pairs of predicted values among all comparable pairs in the test set. The 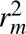 metric is a commonly used performance metric for quantitative structure-activity relationship (QSAR) prediction models, which penalizes large differences between observed and predicted values. It is defined as 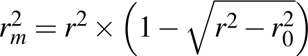 where *r*^2^ and 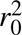 are the squared correlation coefficients between observed and predicted affinities with and without intercept, respectively^47,48^. In addition, we also computed coefficients of determination (*R*^2^; Supplementary Table 3), as *R*^2^ is a widely used measure of quantitative prediction accuracy in the machine learning literature.

ProSmith shows improved overall performance compared to previous methods. On the random split (Table 1), ProSmith exhibits a concordance index of CI = 0.911, which is highly similar but slightly lower compared to the previous state-of-the-art method, NHGNN-DTA^2^ (CI = 0.914) – thus, of all comparable pairs, NHGNN-DTA ranks 0.3% more correctly. However, ProSmith achieves significant improvements over all previous methods in terms of mean squared error (MSE) and the 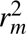 metric. ProSmith is the first method to achieve an MSE below 0.19 on this dataset, lowering the MSE by 0.010 compared to NHGNN-DTA.

**Table 1.**
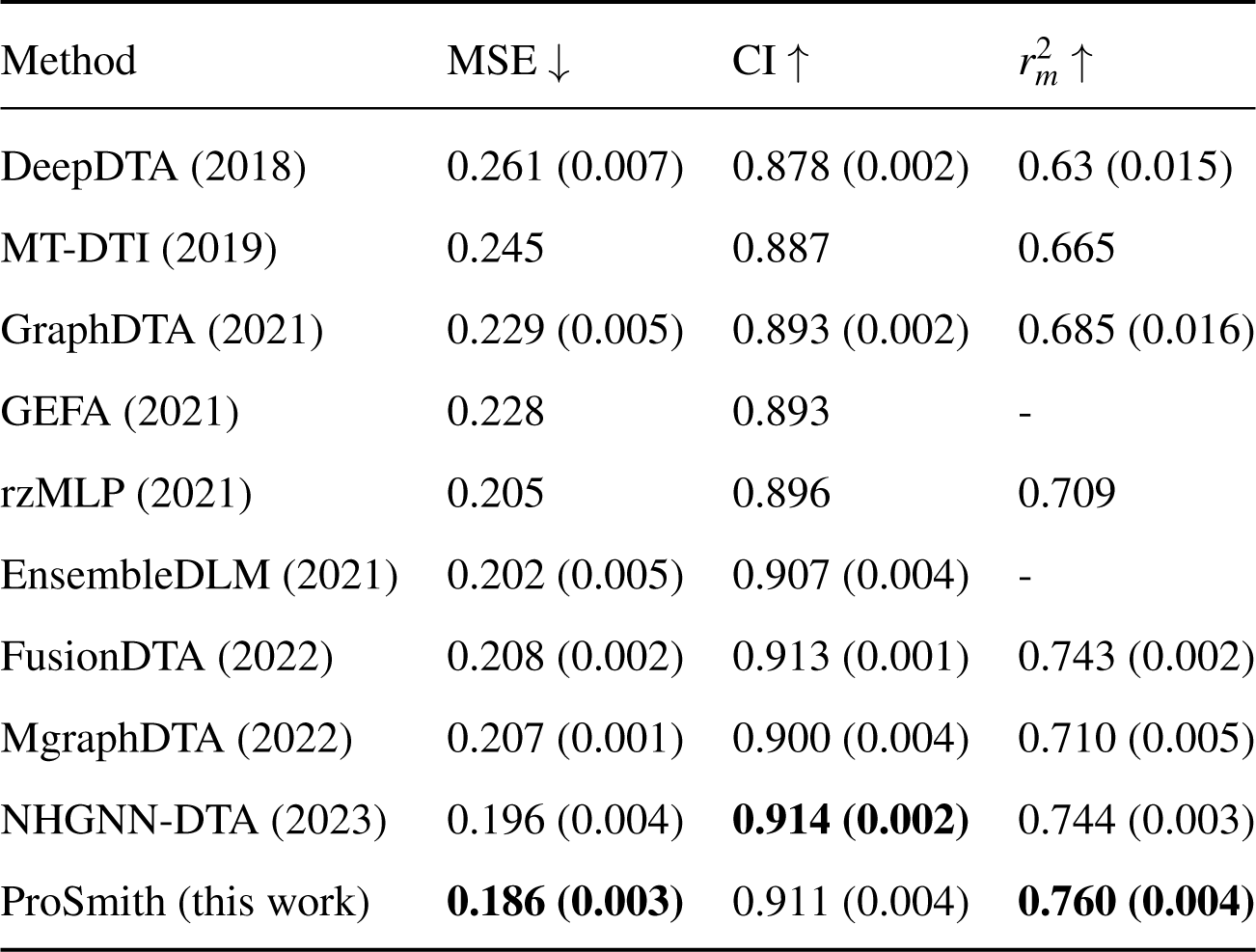
Performance metrics for ProSmith and previously published methods for DTA prediction on the random split of the Davis dataset. Bold numbers highlight the best performance for each metric. Numbers in brackets indicate the standard deviation across the 5 repeated training runs with different splits. Numbers after the method name show year of publication. Performance scores, except for the results of ProSmith, are taken from Ref. [2]. Arrows next to the metric names indicate if higher (*↑*) or lower (*↓*) values correspond to better model performance.

In the more practically relevant scenarios where the drug and/or target were not included in the training set, ProSmith also outperforms all previous methods in almost all comparisons (Table 2). In the two scenarios that exclusively contain target proteins not present in the training data (cold target and cold drug & target), ProSmith achieves substantial performance improvements, clearly surpassing all previous models across all three performance metrics. In the cold drug scenario, ProSmith achieves comparable but slightly worse MSE (0.578 vs. 0.554) and CI (0.733 vs. 0.752) values compared to some previous methods.

**Table 2.** Performance metrics for ProSmith and previously published methods for DTA prediction for different splitting scenarios. Bold numbers highlight the best performance for each metric under each scenario. Numbers in brackets indicate the standard deviation across the 5 repeated training runs with different splits. Arrows next to the metric names indicate if higher (*↑*) or lower (*↓*) values correspond to better model performance. Performance scores, except for the results of ProSmith, are taken from Ref.^2^. In contrast, ProSmith demonstrates a clear improvement in 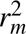 (0.225 compared to the best previous score

| Scenario | Method | MSE ↓ | CI ↑ | $r_m^2$ ↑ |
| --- | --- | --- | --- | --- |
| Cold target | GraphDTA | 0.510 (0.086) | 0.729 (0.012) | 0.154 (0.014) |
|  | GEFA | 0.433 (0.022) | 0.759 (0.009) | 0.289 (0.016) |
|  | FusionDTA | 0.364 (0.021) | 0.826 (0.011) | 0.435 (0.023) |
|  | MgraphDTA | 0.359 (0.023) | 0.813 (0.008) | 0.425 (0.028) |
|  | NHGNN-DTA | 0.344 (0.029) | 0.855 (0.016) | 0.479 (0.021) |
|  | ProSmith (this work) | **0.294 (0.048)** | **0.870 (0.016)** | **0.602 (0.057)** |
| Cold drug | GraphDTA | 0.920 (0.029) | 0.678 (0.036) | 0.160 (0.019) |
|  | GEFA | 0.847 (0.012) | 0.709 (0.028) | 0.182 (0.015) |
|  | FusionDTA | 0.581 (0.094) | 0.737 (0.012) | 0.187 (0.034) |
|  | MgraphDTA | 0.563 (0.065) | 0.729 (0.022) | 0.192 (0.021) |
|  | NHGNN-DTA | **0.554 (0.091)** | **0.752 (0.017)** | 0.207 (0.030) |
|  | ProSmith (this work) | 0.578 (0.006) | 0.733 (0.027) | **0.225 (0.054)** |
| Cold drug and target | GraphDTA | 0.968 (0.096) | 0.579 (0.017) | 0.026 (0.016) |
|  | GEFA | 0.944 (0.092) | 0.610 (0.029) | 0.032 (0.022) |
|  | FusionDTA | 0.876 (0.091) | 0.645 (0.043) | 0.072 (0.048) |
|  | MgraphDTA | 0.874 (0.090) | 0.636 (0.021) | 0.071 (0.041) |
|  | NHGNN-DTA | 0.857 (0.096) | 0.665 (0.038) | 0.087 (0.051) |
|  | ProSmith (this work) | **0.663 (0.159)** | **0.672 (0.005)** | **0.148 (0.097)** |

We assessed the predictive capabilities of ProSmith for drugs with different occurrence frequencies in the training set. We generated new training, validation, and test splits, varying the presence of the test drugs in the training set. As expected, model performance improves with increasing occurrence frequency of a test drug in the training data (Figure 2a). Accuracy is low for drugs occurring between 0 and 10 times in the training set. High prediction performance appears to require at least 30 drug-target pairs with the same drug in the training data. This observation contrasts with our finding in a previous study for predicting enzyme-substrate pairs that two training data points with a given substrate already facilitate accurate predictions^19^. Thus, it appears that learning drug-target interactions is much more difficult than learning enzyme-substrate relationships. That enzyme-substrate relationships are easier to learn may be related to the evolution of dedicated binding sites in response to natural selection for the binding of specific substrates, leading to recognizable signatures in the amino acid sequence.

**Figure 2.**
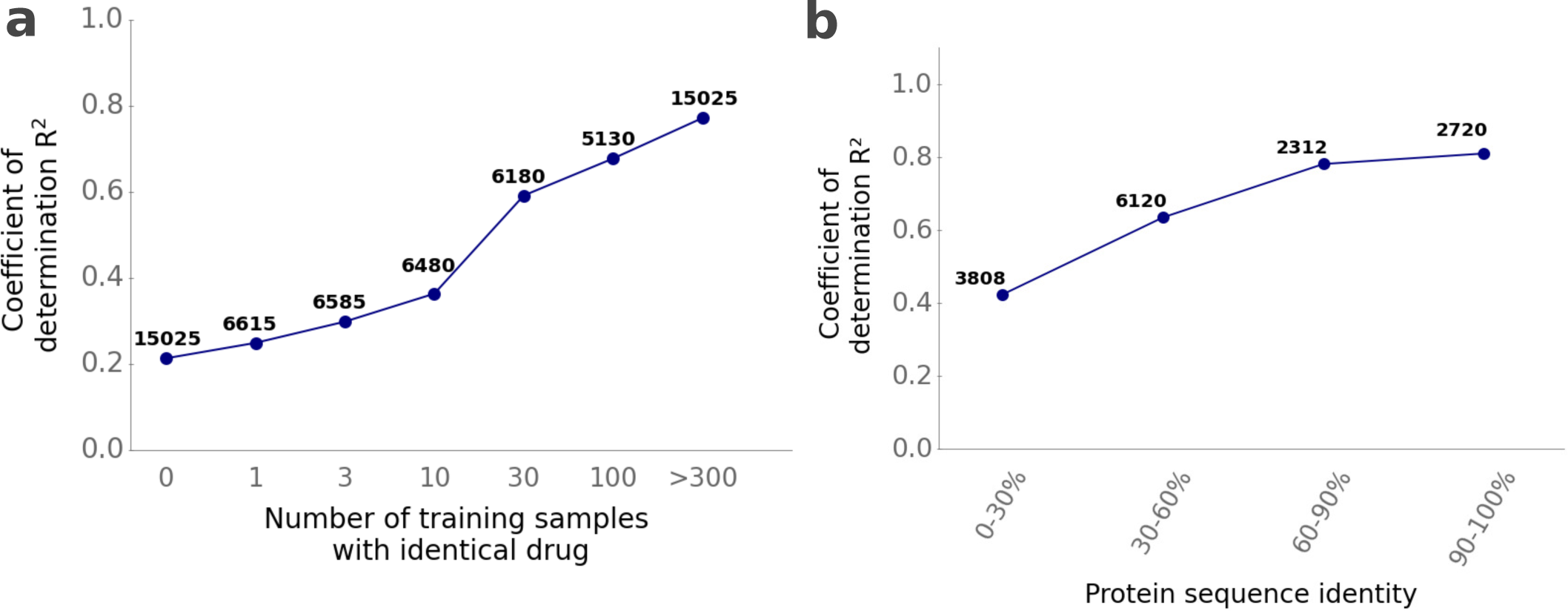
For accurate DTA predictions, ProSmith requires training on identical drugs but not on similar proteins. **(a)** We separately analyzed model performance for dataset splits where drugs from the test set occur in the training set for a specified number of times (0, 1, 3, 10, 30, 100, and > 300). We calculated the coefficient of determination *R*^2^ for each of those test sets separately. **(b)** We divided all five randomly created test sets under the cold target splitting scenario into subsets with different levels of protein sequence identity compared to proteins in the training set, calculating the coefficient of determination *R*^2^ for each subset separately. Numbers above the plotted points indicate the number of test data points in each category. of 0.207). It is important to note that previous methods did not provide the exact training-validation-test splits, and thus, model performances were not evaluated using the exact same test data. However, since all performance scores result from randomly repeating the same splitting procedure five times, the comparison remains meaningful.

We also examined how model performance was influenced by the maximal sequence identity of a test target protein compared to proteins in the training set. This analysis is possible using the cold target data. As expected, increasing protein similarity to training data again leads to improved predictions (Figure 2b). Notably, ProSmith achieves favorable results even for target proteins that are at most very distantly related to any proteins in the training set (maximal sequence identity *<* 30%), explaining over 40% of the variance in the target variable. Overall, our findings highlight the superior performance of ProSmith for DTA predictions compared to previous approaches. The results again highlight the model’s remarkable ability to generalize to previously unseen proteins.

### ProSmith leads to improved generalization for enzyme-substrate pair prediction

Arguably the most comprehensive high-quality resource of protein sequence and functional information is UniProt^49^. While this database lists over 36 million enzymes, less than 1% of these entries contain high-quality annotations of the catalyzed reactions. Thus, the functions of more than 99% of putative enzymes are currently unknown. To address this challenge, we previously developed ESP, a method that predicts whether a small molecule is a potential substrate for a given enzyme based on the enzyme amino acid sequence and on structural information for the small molecule^19^. The ESP gradient boosting model uses as inputs an enzyme representation from the ESM-1b Transformer model – after task-specific fine-tuning – and a small molecule representation generated through a Graph Neural Network (GNN). ESP, which is currently the only general model for the prediction of enzyme-substrate pairs, achieves an accuracy of over 91% on this binary classification task. However, the ESP model fails to produce reliable predictions for small molecules that occurred in the training set only once or not at all.

Here, we use the training and test data from the ESP study to train and test ProSmith for the same enzyme-substrate prediction task. The ESP datasets^19^ consist of positive enzyme-substrate pairs with experimental evidence, complemented with sampled negative enzyme-small molecule pairs, with a positive-to-negative ratio of 1 to 3. The training set comprises 55 418 training data points, while the test set contains 13 336 data points. To perform hyperparameter optimization, we further partitioned the training set into 90% training data and 10% validation data.

Given the requirement for a substantial number of data points for training the multimodal Transformer Network, we expanded the training data – but not the test data – by including data with phylogenetic evidence in addition to the data with experimental evidence. This training set, which comprises a total of 850 291 data points, was already utilized in the ESP study to fine-tune the ESM-1b Transformer Network. To reduce the training time associated with this large training set for the multimodal Transformer Network, we increased the batch size from 12 to 24 for this particular task. While we trained the Transformer Network with the expanded training set, we subsequently trained the gradient boosting models using only the smaller training set based on experimental evidence.

The ProSmith results show remarkable improvements over the original ESP model (Table 3). Notably, the accuracy increased from 91.5% to 94.2%, the ROC-AUC score increased from 0.956 to 0.972, and the Matthews correlation coefficient (MCC), which measures correlation in binary data^50^, increased from 0.78 to 0.85. With these results, ProSmith narrows the gap between the performance of the best available method and perfect predictions by over 30% across all three performance metrics.

**Table 3.** Performance metrics for ProSmith and ESP for the prediction of enzyme-substrate pairs. Bold numbers highlight the best performance for each metric. Arrows next to the metric names (*↑*) indicate that higher values correspond to better model performance.

| Method | Accuracy ↑ | MCC ↑ | ROC-AUC ↑ |
| --- | --- | --- | --- |
| ESP | 91.5% | 0.78 | 0.956 |
| ProSmith | **94.2%** | **0.85** | **0.972** |

A key advancement achieved by ProSmith lies in its ability to produce reliable predictions for small molecules that are not represented multiple times in the training set. As shown in Figure 3a, ProSmith (blue dots) increases the MCC from 0.00 to 0.29 for small molecules not present in the training set and from 0.28 to 0.69 for those present only once (Figure 3a). Furthermore, we investigated the predictive capabilities of ProSmith for enzymes that exhibit different levels of sequence similarity compared to proteins in the training set. Mirroring the results for small molecules, ProSmith leads to the most significant improvements for enzymes dissimilar to any protein in the training set (Figure 3b): for test enzymes with less than 40% sequence identity to any protein used for training, the MCC improves from 0.70 to 0.78. In sum, ProSmith clearly surpasses the performance of the original ESP model, showing a much better ability to generalize to small molecules and enzymes with limited representation in the training set.

**Figure 3.**
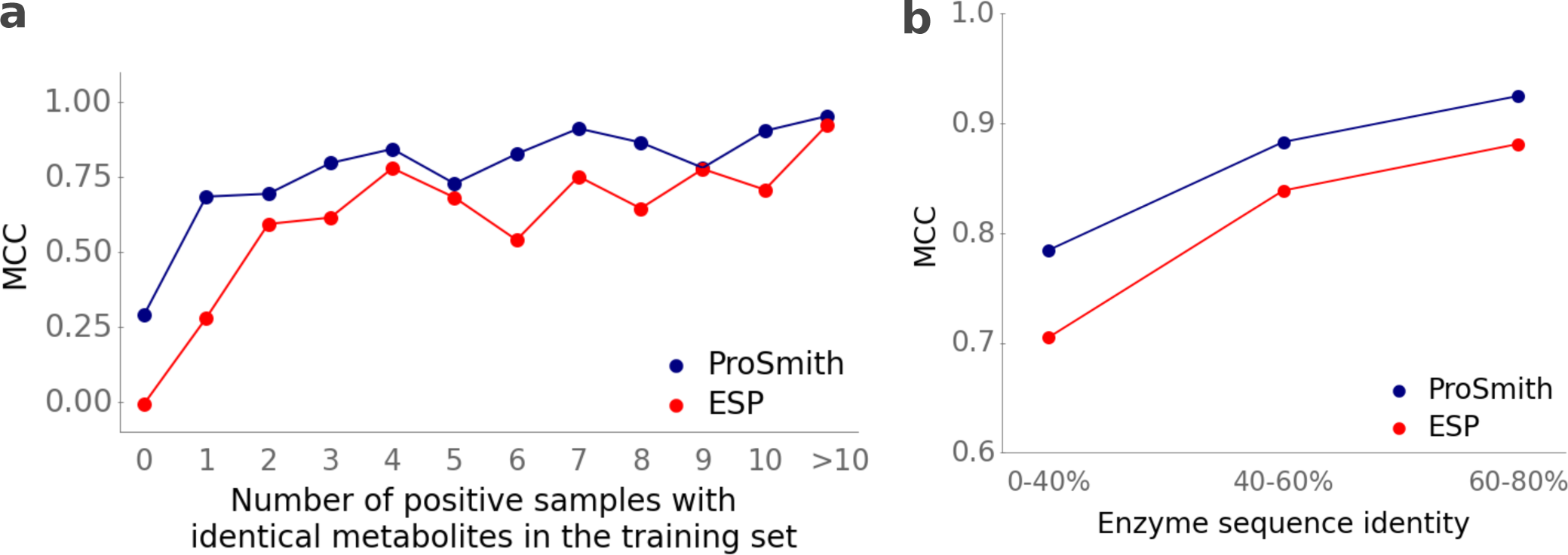
ProSmith outperforms the ESP model in the prediction of enzyme-substrate pairs especially for molecules with limited representation in the test data. **(a)** We grouped small molecules from the test set by how often they occur as substrates among all positive data points in the training set, calculating the MCC for each group separately. **(b)** We divided the test set into subsets with different levels of maximal enzyme sequence identity compared to enzymes in the training set, calculating the MCC for each group separately. The numbers of data points within each subset are listed in Supplementary Tables 4 and 5.

### ProSmith facilitates improved predictions for enzyme-substrate affinities

The third protein-small molecule interaction task that we investigated is predicting the Michaelis constants *K*_M_ of enzyme-substrate pairs. *K*_M_ represents the substrate concentration at which an enzyme operates at half of its maximal catalytic rate, and thus indicates the affinity of an enzyme for a specific substrate. Knowledge of *K*_M_ values is crucial for understanding enzymatic interactions between enzymes and metabolites, as it relates the intracellular concentration of a metabolite to its consumption rate.

For this task, we utilize a dataset containing 11 676 experimental *K*_M_ measurements, which we had compiled to develop a previous model that predicts Michaelis constants^29^. We adopted the same split as used in that study, which divides the *K*_M_ dataset into 80% training data and 20% test data. To obtain a validation set, we further split the original training set into 10% validation data and 90% training data.

Similar to the situation encountered for the DTA prediction task, the number of available *K*_M_ data points is relatively small for training the ProSmith Transformer Network. We thus used the enzyme-substrate prediction task for pre-training, i.e., we initialized the ProSmith Transformer Network for *K*_M_ with the final parameters from training the model for the enzyme-substrate prediction. This initialization provides a starting point that allows the model to leverage previously learned knowledge. We also tested using the model parameters that resulted from pre-training the ProSmith Transformer Network on the IC50 values, which we used above for the DTA model. However, this led to slightly worse results.

ProSmith demonstrates superior performance compared to two previous *K*_M_ prediction models that utilized the same training and test data^28,29^ (Table 4). Similar to what was seen for the other two prediction tasks, ProSmith enhances the ability to generalize to proteins that differ significantly from those in the training set (Supplementary Figure 1a). However, its capacity to generalize to unseen substrates remains limited and is very similar to the previous state-of-the-art method (Supplementary Figure 1b).

**Table 4.** Performance metrics of ProSmith and previously published methods for the prediction of Michaelis constants. *K*_M_. Metrics were calculated using the same training and test data for all three models. Bold numbers highlight the best performance for each metric. Arrows next to the metric names indicate if higher (*↑*) or lower (*↓*) values correspond to better model performance.

| Method | MSE ↓ | $R^2$ ↑ | Pearson $r$ ↑ |
| --- | --- | --- | --- |
| ENKIE (2022) | - | 0.463 | 0.680 |
| Kroll et al. (2021) | 0.653 | 0.527 | 0.728 |
| ProSmith (this work) | **0.604** | **0.563** | **0.752** |

Although the overall model performance exhibits clear improvement, the magnitude of performance gain is smaller compared to the enzyme-substrate prediction and DTA prediction tasks. We hypothesize that this comparatively small improvement may be related to the relatively low number of training data points in comparison to the other two tasks. We tentatively conclude that ProSmith yields promising results even with small datasets, but its greatest performance gains are observed when applied to larger training datasets.

### Final predictions use complementary information from the multimodal Transformer Network and the protein/small molecule representations

To obtain its final predictions, ProSmith calculates a weighted mean across the results of three distinct gradient boosting models with different inputs. The weights for the weighted mean calculation are treated as hyperparameters, i.e., they are chosen such that they maximize the performance on the validation set for a given task. Figure 4 shows the weights assigned to each model for all investigated prediction tasks; for the DTA prediction, we calculated the mean across all 5 random splits for each splitting scenario.

**Figure 4.**
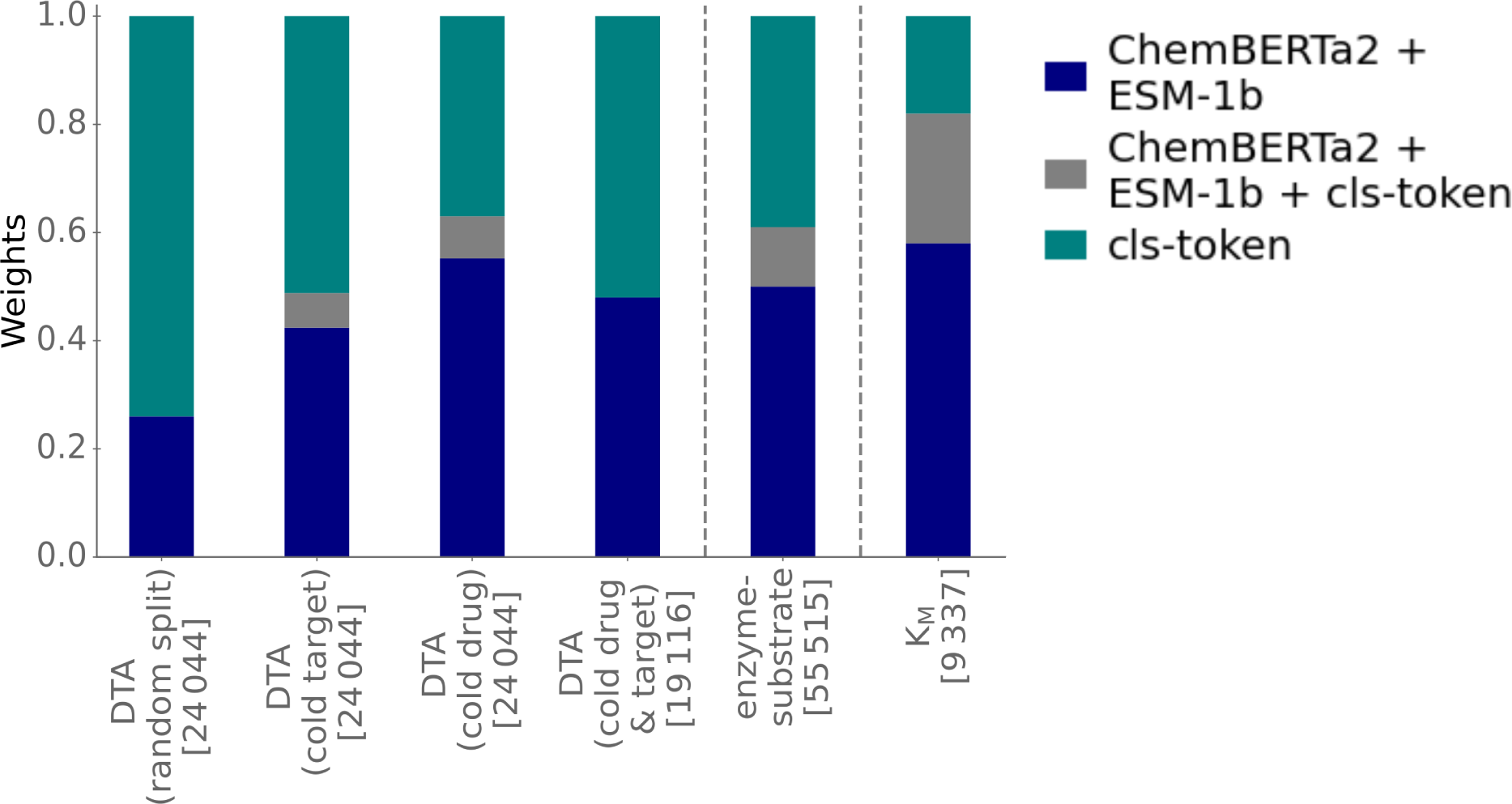
The optimal ProSmith models combine predictions based on the multimodal Transformer Network with predictions based on separate numerical representations of proteins and small molecules. The bar plots quantify the weights assigned to the predictions of the three distinct gradient boosting models contributing to ProSmith: the model trained only on the cls token from ProSmith’s multimodal Transformer Network (teal); the model combining ESM-1b and ChemBERTa2 vectors (blue); and the model combining all three input vectors (grey). The weights are displayed separately for the distinct prediction tasks: drug-target affinity (DTA) (four different splits); enzyme-substrate pairs; Michaelis constants *K*_M_. Numbers in square brackets show the number of training data points.

The model using solely the cls token as its input and the model combining the ChemBERTa2 and ESM-1b vectors as its inputs have the greatest influence on the final predictions (Figure 4). It is likely that some relevant information from the 1280-dimensional ESM-1b vector and the 600-dimensional ChemBERTa2 vector cannot be captured fully within the 768-dimensional cls token, which also stores information about the protein-small molecule interaction. Adding separate, general protein and small molecule information appears to be advantageous for the generation of more accurate overall predictions. Our findings indicate that the combination of multiple gradient boosting models trained on different input information yields better and more robust performance compared to a single model utilizing all input information (Supplementary Table 6), consistent with previous observations^30^.

When predicting Michaelis constants *K*_M_, the model utilizing only the cls token exhibits the lowest influence on model performance (Figure 4). This observation might be related to the limited number of only ∼ 9000 data points available for training the ProSmith Transformer Network for this task. For tasks with more extensive training data, the influence of the cls token on model predictions is much more substantial. These findings indicate that the ProSmith model can adapt its predictions based on the availability of training data points, optimizing model performance accordingly.

## Discussion

In this study, we introduce ProSmith, a novel machine learning framework for predicting the interactions between proteins and drugs or other small molecules. The main methodological advance of our model is the utilization of a multimodal Transformer Network that can effectively process amino acid sequences of proteins and SMILES string representations of small molecules within the same input sequence (Figure 1). The ability of this model architecture to incorporate information on the interaction between a protein and a small molecule during the generation of the corresponding numerical representation leads to a superior ability of the trained model to predict drug-target interactions for target proteins dissimilar to those included in the training data.

We demonstrate that the proposed ProSmith framework does not only lead to improved performance on drug-target interaction predictions, but also on other protein-small molecule interaction tasks. Thus, ProSmith outperforms previous state-of-the-art methods in predicting protein-small molecule interactions across diverse tasks of high relevance to biomedical, biotechnological, and biological research. Our results highlight the potential of leveraging multimodal inputs to achieve significant advancements in predicting complex molecular interactions. The proposed framework is not limited to modeling protein-small molecule interactions. For example, a similar approach could be employed to predict protein-reaction interactions, which would be useful for the prediction of enzymatic turnover numbers *k_cat_*^30,31^.

Due to the computational expenses associated with training Transformer Networks, we did not fully optimize ProSmith’s performance for each individual task. Instead we chose the hyperparameters through trial and error for the drug-target interaction predictions, and we used the same Transformer Network hyperparameters for the two additional tasks. However, hyperparameter search for the Transformer Network, such as optimizing the learning rate, batch size, number of layers, and embedding dimensions, is crucial for improving model performance. In particular, while recent research suggests that some capabilities of Transformer Networks only emerge after surpassing a certain network size limit^51^, limited computational resources led us to choose only six transformer layers. An extensive hyperparameter search, executed separately for each individual task, will likely lead to more suitable ProSmith model architectures with improved results.

For the token embeddings of protein amino acid sequences and SMILES strings, we utilized pre-trained embeddings provided by the protein Transformer Network ESM-1b^13^ and the SMILES Transformer Network ChemBERTa2^14^. The parameters of these two networks were not adjusted during the training of ProSmith. In future investigations, it would be valuable to also explore the impact of adjusting the weights of these embeddings simultaneously with the weights of the ProSmith Transformer Network.

Ensemble modeling has been proven effective in enhancing DTA prediction models, as demonstrated in a previous study^9^. Averaging the predictions of multiple well-performing models, including ProSmith, could yield further performance gains. For instance, while ProSmith exhibits a slightly lower concordance index (CI) than the previous state-of-the-art method NHGNN-DTA^2^ in the random splitting scenario, the latter shows worse MSE and *r*^2^ metric scores. Combining these models through a weighted mean prediction approach is likely to overcome the limitations of the individual models, achieving state-of-the-art performance across all three evaluation metrics.

ProSmith appears to show the most substantial performance gains when trained on larger datasets. In deep learning, it is common to pre-train models on similar tasks with more abundant data when training data is limited^44,45^. We did this for two of the task explored above, pre-training ProSmith on IC50 values before training the DTA prediction model, and on the enzyme-substrate pair data before training the *K*_M_ prediction model. Previous studies have shown the benefits of training only the last layers of pre-trained models while keeping the initial layers fixed^45,52^. Investigating the applicability of this approach to protein-small molecule interaction tasks with small training datasets, such as for the *K*_M_ prediction, could be another avenue for future exploration.

The drug-target affinity prediction model can be retrained by researchers interested in the interaction of alternative drug candidates with potential targets using their own data. Users can employ the Python functions provided on GitHub (https://github.com/AlexanderKroll/ProSmith) to train the ProSmith model for arbitrary protein-small molecule interaction tasks on datasets of up to ∼100,000 data points within a reasonable time frame and without the requirement of an extensive GPU infrastructure, as detailed in Supplementary Text.

## Methods

### Software and code availability

All software was coded in Python^53^. We implemented the multimodal Transformer Network in PyTorch^54^. We fitted the gradient boosting models using the library XGBoost^38^. The code used to generate the results of this paper is available from https://github.com/AlexanderKroll/ProSmith in the form of Jupyter notebooks. All datasets used to create the results of this manuscript are available from https://doi.org/10.5281/zenodo.8182031.

### Calculation of protein token embeddings

We use protein amino acid sequences to represent proteins in the input of the ProSmith Transformer model. Every amino acid in a sequence is represented through a separate token. To numerically encode information about the token, we used learned representations from the ESM-1b model, a Transformer Network with 33 layers that was trained on ∼27 million protein sequences^13^. We applied the trained ESM-1b model to each protein amino acid sequence and extracted the updated 1280-dimensional token representations from the last layer of the model.

### Calculation of small molecule token embeddings

We used SMILES strings to represent small molecules in the input of the ProSmith Transformer. To divide the SMILES string into disjoint tokens, we used the ChemBERTa2 model^14^. ChemBERTa2 is a Transformer Network with 3 layers that was trained on ∼77 million different SMILES strings. We applied this model to each SMILES string in our dataset, and we extracted 600-dimensional learned token embeddings from the last layer of ChemBERTa2.

### Input representation of the multimodal Transformer Network

Every input sequence of the multimodal Transformer Network has the following structure: first the ‘cls’ classification token, then the protein amino acid sequence tokens, followed by a separation token, and finally, the SMILES string tokens. In the input, the cls token is represented by a vector of all ones, the separation token is a vector of all zeros, and the protein and SMILES tokens were extracted from ESM-1b and ChemBERTa2, respectively, as described above. The maximum length for protein sequences was set to 1024 and the maximum number of tokens for SMILES strings was set to 256. For longer amino acid sequences, we only kept the first 1024 amino acids; for longer smiles strings, we kept only the fist 256 tokens.

### Model architecture of the multimodal Transformer Network

Before being processed by the multimodal Transformer Network, the amino acid tokens are fed through a protein pooling layer, and the SMILES tokens are fed through a SMILES pooling layer. Each of the two pooling layers is a single-layered fully connected neural network with the ReLU activation function. The pooling layers are applied to each token embedding, mapping the embeddings to the hidden dimension of our multimodal Transformer Network, 768. The parameters of the pooling layers are updated in each iteration of training the Transformer Network.

The classification, protein, separation, and SMILES token embeddings of dimension 768 are used as the input of a Transformer Network called BERT, which stands for Bidirectional Encoder Representations from Transformers^45^. The number of Transformer Layers was set to 6, each with 6 attention heads. The activation function was set to GELU, which is a smoothed version of the ReLU activation function.

After updating each token in the input sequence six times, we extract the updated 768-dimensional representation of the classification token and pass it through a fully connected neural network with one hidden layer of dimension 32 and ReLU as the activation function. The output layer has one node; it uses no activation function for regression tasks and the sigmoid activation function for binary classification tasks.

### Training of the multimodal Transformer Network

We trained the whole model described above end-to-end, i.e., the BERT model together with the pooling layers and the fully connected layers applied to the update classification token. The learning rate was set to 10*^−^*^5^. The loss function was set to the mean squared error for regression tasks and to the binary cross entropy for binary classification tasks. We trained each Transformer Network for 100 epochs and saved model parameters after each epoch. After training, to guard against overfitting, we selected the model that achieved the best performance on the validation set.

### Processing of batches for the Transformer Network training

Storing all protein sequence tokens and all SMILES string tokens during training requires too much RAM for large datasets. To overcome this issue, we divided the set of all proteins into smaller subsets of size 1000, and we did the same for the set of all SMILES strings. During training, we only load one subset of protein sequences tokens and one subset of SMILES sequence tokens at a time into the RAM, and we iterate over all possible combinations of protein and SMILES subset combinations.

### pre-training of the Transformer Network on the IC50 dataset

We downloaded the Ligand-Target-Affinity Dataset from BindingDB^46^. We extracted all drug-target pairs with experimentally measured IC50 values from this dataset. We excluded all pairs where either the drug or the target were present in the Davis dataset^43^. This resulted in a dataset with 1 039 565 entries. We split this dataset into 95% training data and 5% validation data. We used the training data to pre-train the ProSmith Transformer Network for the drug-target affinity (DTA) task. We trained the Transformer Network for 100 epochs and saved model parameters after each epoch. After training, we selected the model that achieved the best performance on the validation set. Because of the large training set size, we used a higher batch size of 192 compared to the other tasks investigated in this study. As is common for larger batch sizes, we also increased the learning rate slightly to 1.5 × 10^−5^.

### Splitting the Davis dataset

The Davis dataset consists of 30 056 data points with 72 different drugs and 442 proteins with measured *K_d_* values. To split this dataset into training, validation, and test sets, we adopted the identical strategy employed by the previously leading method, NHGNN-DTA^2^. We generated five random splits for each of four scenarios: random; cold target; cold drug; and cold drug & target (for details, see the section “ProSmith leads to improved generalization for drug-target affinity predictions” in the Results section).

### Splitting Davis data with different occurrence frequencies of test drugs in the training set

To assess the predictive capabilities of ProSmith for drugs with different occurrence frequencies in the training set, we generated new dataset splits. We split the data in such a way that for 15 randomly selected drugs from the test set, only 1, 3, 10, 30, or 100 drug-target pairs with the same drug but paired with different targets are present in the training set. Model performance for drugs that do not occur in the training set or that are present more than 300 times was extracted from the results for the cold drug and random splitting scenario, respectively.

### Training of the gradient boosting models

To find the best hyperparameters for the gradient boosting models, we performed a random grid search with 2 000 iterations. In each iteration, we trained a gradient boosting model with a different set of hyperparameters on the training data and assessed the performance of the resulting model on the validation set. After this random search, we selected the hyperparameter set that led to the best performance on the validation set. We used the Python package hyperopt^55^ to perform the hyperparameter optimization for the following hyperparameters: learning rate, maximum tree depth, lambda and alpha coefficients for regularization, maximum delta step, minimum child weight, and number of training epochs. For the task of predicting enzyme-substrate pairs, we added a weight for the negative data points. This hyperparameter was added because the dataset is imbalanced, and it allows the model to assign a lower weight to the overrepresented negative data points during training. We used the Python package xgboost^38^ for training the gradient boosting models.

### Computational resources

To train the Transformer Networks and to perform hyperparameter optimization for all gradient boosting models, we used the High Performance Computing Cluster at Heinrich Heine University Düsseldorf (Germany). All training processes were executed on a single NVIDIA A100 GPU. The only exception was the pre-training of the Transformer Network for the IC50 value prediction. To shorten the training time for the large training set with ∼ 1 million data points, we trained this model on four NVIDIA A100 GPUs.

## Acknowledgements

We thank Martin K. M. Engqvist for interesting discussions on multimodal Transformer Networks. Computational support and infrastructure was provided by the “Centre for Information and Media Technology” (ZIM) at Heinrich Heine University Düsseldorf, Germany. This work was funded through grants to MJL by the European Union (ERC AdG “MechSys”–Project ID 101055141) and by the Deutsche Forschungsgemeinschaft (DFG, German Research Foundation: CRC 1310, and, under Germany’s Excellence Strategy, EXC 2048/1–Project ID 390686111).

## Author contributions

AK conceived of the study, performed all analyses, and drafted the manuscript. MJL supervised the study and acquired funding. SR and AK designed and implemented the multimodal Transformer Network. MJL and AK interpreted the results and edited the manuscript.

## Supplementary Information

### Supplementary Text

#### The ProSmith Transformer Network can be trained with limited computational resources

The training of Transformer Networks typically demands extensive computational resources. For example, the ESM-1b Transformer Network, designed for protein amino acid sequences, was trained on a compu-tation cluster with 64 individual NVIDIA V100 GPUs, each equipped with 16 GB RAM, over a period of approximately 19 days. In contrast, because of its comparatively small size, we were able to train the ProSmith Transformer Network using a single NVIDIA A100 GPU with 40 GB RAM in a much shorter timeframe; training on the Davis dataset for 100 epochs required only ∼22 hours.

In tests on a more affordable GPU – a single NVIDIA RTX6000 with 24GB RAM – we obtained the same model performance with an acceptable increase in training time (∼39 hours). Thus, interested users can train the ProSmith model for arbitrary protein-small molecule interaction tasks on datasets of up to ∼100,000 data points within a reasonable time frame and without the requirement of an extensive GPU infrastructure. Such future applications are greatly facilitated by the user-friendly Python functions provided on our GitHub repository.

## Supplementary Tables

**Supplementary Table 1.** Hyperparameters of the ProSmith Transformer Network.

| Hyperparameter | Value |
| --- | --- |
| learning rate | $10^{-5}$ |
| hidden dimension | 768 |
| number of layers | 6 |
| number of attention heads | 6 |
| batch size | 12 |

**Supplementary Table 2.** Hyperparameters of the ProSmith gradient boosting models.

| Hyperparameter | Search space |
| --- | --- |
| learning rate | 0.01 – 0.5 |
| maximum depth of trees | 6 – 14 |
| regression coefficient alpha | 0 – 5 |
| regression coefficient lambda | 0 – 5 |
| maximum delta step | 0 – 5 |
| minimum child weight | 0.1 – 15 |
| number of trees | 30 – 1000 |

**Supplementary Table 3.** Coefficient of determination *R*^2^ of ProSmith for different splitting scenarios of the Davis dataset. Numbers in brackets indicate the standard deviation for the results of the 5 repeated training runs with different splits.

| Scenario | $R^2$ |
| --- | --- |
| random | 0.772(0.010) |
| cold target | 0.676(0.061) |
| cold drug | 0.213(0.007) |
| cold target and drug | 0.098 (0.129) |

**Supplementary Table 4.** We divided the ESP test set into subsets according to the number of positive data points with the same small molecule in the training set. The table shows the number of test data points in each subset.

| Number of positive samples with identical metabolite in the training set | Size of subset |
| --- | --- |
| 0 | 602 |
| 1 | 495 |
| 2 | 385 |
| 3 | 501 |
| 4 | 384 |
| 5 | 336 |
| 6 | 330 |
| 7 | 329 |
| 8 | 201 |
| 9 | 337 |
| 10 | 126 |
| > 10 | 9265 |

**Supplementary Table 5.** We divided the ESP test set into subsets according to the maximal sequence identity of an enzyme compared to all training enzymes. The table shows the number of test data points in each subset.

| Maximal sequence identity | Size of subset |
| --- | --- |
| 0-40% | 6044 |
| 40-60% | 4050 |
| 60-80% | 3164 |

**Supplementary Table 6.** Performance metrics for all three trained gradient boosting models with different input vectors and for their combined weighted mean prediction. For the test sets for the *K*_M_ prediction task and for the four splitting scenarios of the Davis dataset, the table lists coefficients of determination *R*^2^; for the enzyme-substrate prediction, the table lists Matthews correlation coefficients (MCCs).

|  | ChemBERTa2 + ESM-1b | ChemBERTa2 + ESM-1b + cls | cls token | weighted mean |
| --- | --- | --- | --- | --- |
| Enzyme-substrate prediction | 0.769 | 0.838 | 0.840 | 0.850 |
| $K_M$ prediction | 0.528 | 0.509 | 0.541 | 0.563 |
| Davis (random) | 0.727 | 0.747 | 0.763 | 0.772 |
| Davis (cold target) | 0.623 | 0.662 | 0.658 | 0.676 |
| Davis (cold drug) | 0.127 | 0.115 | 0.095 | 0.204 |
| Davis (cold drug & target) | 0.034 | 0.044 | 0.033 | 0.098 |

## Supplementary Figures

**Supplementary Figure 1.**
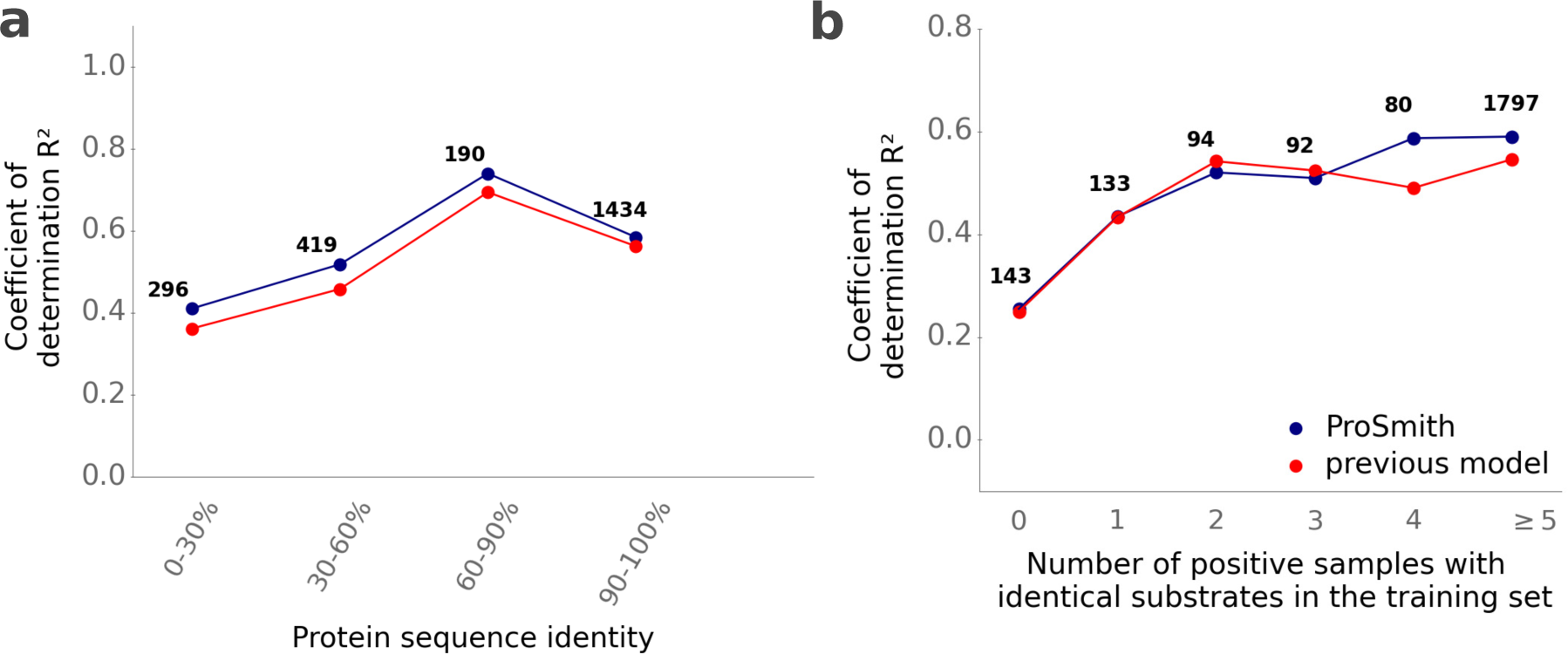
ProSmith outperforms previous models in the prediction of Michaelis constants *K*_M_ especially for enzymes not highly similar to proteins in the training set. **(a)** We divided the test set into subsets with different levels of maximal enzyme sequence identity compared to enzymes in the training set, calculating the MCC for each group separately. **(b)** We grouped substrates from the test set by how often they occur as substrates among all data points in the training set, calculating the MCC for each group separately. Numbers above the plotted points indicate the number of test data points in each category.

## References

1. Wen, M. et al. Deep-learning-based drug–target interaction prediction. J. Proteome Res. 16, 1401–1409 (2017).

2. He, H., Chen, G. & Chen, C. Y.-C. NHGNN-DTA: A Node-adaptive Hybrid Graph Neural Network for Interpretable Drug-target Binding Affinity Prediction. Bioinformatics, btad355 (2023).

3. Yang, Z., Zhong, W., Zhao, L. & Chen, C. Y.-C. ML-DTI: mutual learning mechanism for interpretable drug–target interaction prediction. J. Phys. Chem. Lett. 12, 4247–4261 (2021).

4. Öztürk, H., Özgür, A. & Ozkirimli, E. DeepDTA: deep drug–target binding affinity prediction. Bioinformatics 34, i821–i829 (2018).

5. Shin, B., Park, S., Kang, K. & Ho, J. C. Self-attention based molecule representation for predicting drug-target interaction in Machine Learning for Healthcare Conference (2019), 230–248.

6. Nguyen, T. et al. GraphDTA: Predicting drug–target binding affinity with graph neural networks. Bioinformatics 37, 1140–1147 (2021).

7. Nguyen, T. M., Nguyen, T., Le, T. M. & Tran, T. Gefa: early fusion approach in drug-target affinity prediction. IEEE/ACM Trans. Comput. Biol. Bioinform. 19, 718–728 (2021).

8. Qiu, Z., et al. rzMLP-DTA: gMLP network with ReZero for sequence-based drug-target affinity prediction in 2021 IEEE International Conference on Bioinformatics and Biomedicine (BIBM) (2021), 308–313.

9. Kao, P.-Y., Kao, S.-M., Huang, N.-L. & Lin, Y.-C. Toward drug-target interaction prediction via ensemble modeling and transfer learning in 2021 IEEE International Conference on Bioinformatics and Biomedicine (BIBM) (2021), 2384–2391.

10. Yuan, W., Chen, G. & Chen, C. Y.-C. FusionDTA: attention-based feature polymerizer and knowledge distillation for drug-target binding affinity prediction. Brief. Bioinform. 23, bbab506 (2022).

11. Yang, Z., Zhong, W., Zhao, L. & Chen, C. Y.-C. MGraphDTA: deep multiscale graph neural network for explainable drug–target binding affinity prediction. Chem. Sci. 13, 816–833 (2022).

12. Weininger, D. SMILES, a chemical language and information system. 1. Introduction to methodology and encoding rules. J. Chem. Inf. Comput. Sci. 28, 31–36 (1988).

13. Rives, A. et al. Biological structure and function emerge from scaling unsupervised learning to 250 million protein sequences. PNAS 118, 622226 (2021).

14. Ahmad, W., Simon, E., Chithrananda, S., Grand, G. & Ramsundar, B. Chemberta-2: Towards chemical foundation models. arXiv preprint at arXiv:2209.01712 (2022).

15. Radford, A. et al. Learning Transferable Visual Models From Natural Language Supervision in Proceedings of the 38th International Conference on Machine Learning (eds Meila, M. & Zhang, T.) 139 (PMLR, July 2021), 8748–8763.

16. Alayrac, J.-B., et al. Flamingo: a visual language model for few-shot learning. Adv. Neural Inf. Process. Syst. 35, 23716–23736 (2022).

17. Reed, S., et al. A generalist agent. arXiv preprint at arXiv:2205.06175 (2022).

18. Lin, J., et al. Interbert: Vision-and-language interaction for multi-modal pretraining. arXiv preprint at arXiv:2003.13198 (2020).

19. Kroll, A., Ranjan, S., Engqvist, M. K. & Lercher, M. J. A general model to predict small molecule substrates of enzymes based on machine and deep learning. Nat. Commun. 14, 2787 (2023).

20. Alballa, M., Aplop, F. & Butler, G. TranCEP: Predicting the substrate class of transmembrane transport proteins using compositional, evolutionary, and positional information. PLoS One 15, e0227683 (2020).

21. Mou, Z. et al. Machine learning-based prediction of enzyme substrate scope: Application to bacterial nitrilases. Proteins Struct. Funct. Bioinf 89, 336–347 (2021).

22. Yang, M. et al. Functional and informatics analysis enables glycosyltransferase activity prediction. Nat. Chem. Biol. 14, 1109–1117 (2018).

23. Pertusi, D. A. et al. Predicting novel substrates for enzymes with minimal experimental effort with active learning. Metab. Eng. 44, 171–181 (2017).

24. Röttig, M., Rausch, C. & Kohlbacher, O. Combining structure and sequence information allows automated prediction of substrate specificities within enzyme families. PLoS Comput. Biol. 6, e1000636 (2010).

25. Chevrette, M. G., Aicheler, F., Kohlbacher, O., Currie, C. R. & Medema, M. H. SANDPUMA: ensemble predictions of nonribosomal peptide chemistry reveal biosynthetic diversity across Actinobacteria. Bioinformatics 33, 3202–3210 (2017).

26. Goldman, S., Das, R., Yang, K. K. & Coley, C. W. Machine learning modeling of family wide enzyme-substrate specificity screens. PLoS Comput. Biol. 18, e1009853 (2022).

27. Visani, G. M., Hughes, M. C. & Hassoun, S. Enzyme promiscuity prediction using hierarchy-informed multi-label classification. Bioinformatics 37, 2017–2024 (2021).

28. Gollub, M. G., Backes, T., Kaltenbach, H.-M. & Stelling, J. ENKIE: A package for predicting enzyme kinetic parameter values and their uncertainties. bioRxiv preprint at 10.1101/2023.03.08.531697, 2023–03 (2023).

29. Kroll, A., Engqvist, M. K., Heckmann, D. & Lercher, M. J. Deep learning allows genome-scale prediction of Michaelis constants from structural features. PLoS Biol. 19, e3001402 (2021).

30. Kroll, A., Rousset, Y., Hu, X.-P., Liebrand, N. A. & Lercher, M. J. Turnover number predictions for kinetically uncharacterized enzymes using machine and deep learning. Nat. Commun. 14, 4139 (2023).

31. Li, F. et al. Deep learning-based kcat prediction enables improved enzyme-constrained model reconstruction. Nat. Catal., 1–11 (2022).

32. Borger, S., Liebermeister, W. & Klipp, E. Prediction of enzyme kinetic parameters based on statistical learning. Genom. Inform. 17, 80–87 (2006).

33. Xu, P., Zhu, X. & Clifton, D. A. Multimodal learning with transformers: A survey. IEEE Trans. Pattern Anal. Mach. (2023).

34. Bahdanau, D., Cho, K. & Bengio, Y. Neural machine translation by jointly learning to align and translate. Preprint at 10.48550/arXiv.1409.0473 (2014).

35. Shao, T., Guo, Y., Chen, H. & Hao, Z. Transformer-based neural network for answer selection in question answering. IEEE Access 7, 26146–26156 (2019).

36. Minixhofer, B., Gritta, M. & Iacobacci, I. Enhancing Transformers with Gradient Boosted Decision Trees for NLI Fine-Tuning. arXiv preprint at arXiv:2105.03791 (2021).

37. Friedman, J. H. The elements of statistical learning: Data mining, inference, and prediction (SpringerOpen, 2017).

38. Chen, T. & Guestrin, C. Xgboost: A scalable tree boosting system in Proceedings of the 22nd acm sigkdd international conference on knowledge discovery and data mining (2016), 785–794.

39. Zhou, Z.-H. & Zhou, Z.-H. Ensemble learning (Springer, 2021).

40. Bergstra, J. & Bengio, Y. Random search for hyper-parameter optimization. J. Mach. Learn. Res. 13, 281–305 (2012).

41. Bull, S. C. & Doig, A. J. Properties of protein drug target classes. PLoS One 10, e0117955 (2015).

42. Zhavoronkov, A., Vanhaelen, Q. & Oprea, T. I. Will artificial intelligence for drug discovery impact clinical pharmacology? Clin. Pharmacol. Ther. 107, 780–785 (2020).

43. Davis, M. I. et al. Comprehensive analysis of kinase inhibitor selectivity. Nat. Biotechnol. 29, 1046– 1051 (2011).

44. Krizhevsky, A., Sutskever, I. & Hinton, G. E. Imagenet classification with deep convolutional neural networks. Adv. Neural Inf. Process. Syst. 25 (2012).

45. Devlin, J., Chang, M.-W., Lee, K. & Toutanova, K. Bert: Pre-training of deep bidirectional trans-formers for language understanding. arXiv preprint at arXiv:1810.04805 (2018).

46. Gilson, M. K. et al. BindingDB in 2015: a public database for medicinal chemistry, computational chemistry and systems pharmacology. Nucleic Acids Res. 44, D1045–D1053 (2016).

47. Pratim Roy, P., Paul, S., Mitra, I. & Roy, K. On two novel parameters for validation of predictive QSAR models. Molecules 14, 1660–1701 (2009).

48. Roy, K. et al. Some case studies on application of “rm2” metrics for judging quality of quantitative structure–activity relationship predictions: emphasis on scaling of response data. J. Comput. Chem. 34, 1071–1082 (2013).

49. Consortium, T. U. UniProt: the universal protein knowledgebase in 2021. Nucleic Acids Res. 49, D480–D489 (2020).

50. Chicco, D. & Jurman, G. The advantages of the Matthews correlation coefficient (MCC) over F1 score and accuracy in binary classification evaluation. BMC Genom 21, 1–13 (2020).

51. Srivastava, A., et al. Beyond the imitation game: Quantifying and extrapolating the capabilities of language models. arXiv preprint at arXiv:2206.04615 (2022).

52. Simonyan, K. & Zisserman, A. Very deep convolutional networks for large-scale image recognition. arXiv preprint at arXiv:1409.1556 (2014).

53. Van Rossum, G. & Drake, F. L. Python 3 Reference Manual (CreateSpace, Scotts Valley, 2009).

54. Paszke, A. et al. Pytorch: An imperative style, high-performance deep learning library. Adv. Neur. In. 32, 8026–8037 (2019).

55. Bergstra, J., Yamins, D. & Cox, D. Making a science of model search: Hyperparameter optimization in hundreds of dimensions for vision architectures in International conference on machine learning (2013), 115–123.

